## Supplementary material for "Myristoylated Neuronal Calcium Sensor-1 captures the ciliary vesicle at distal appendages": Source Data 6-uncropped images of the Immunoblot with label

Figure 1D_GFP

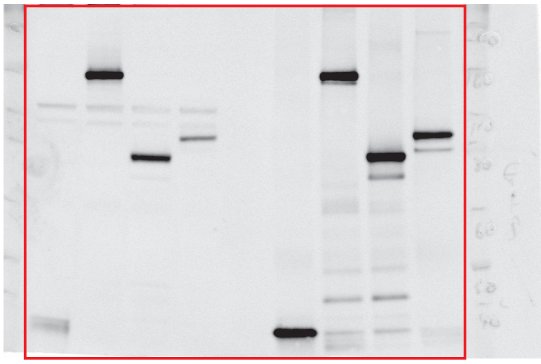

Figure 1D_NCS1

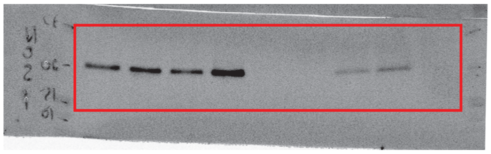

Figure 1D_Tubulin

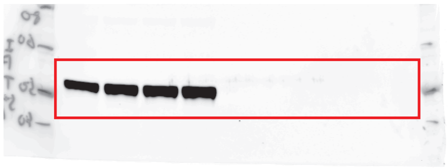

Figure 1E_anti-MYC

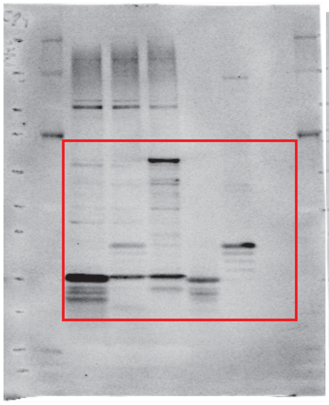

Figure 1E_anti-HA

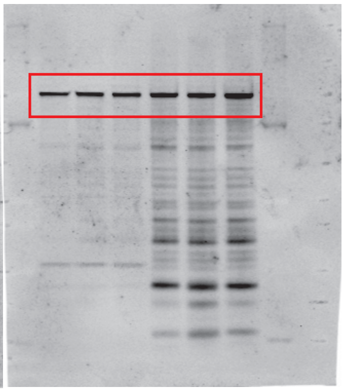

Figure 1F_anti-MYC

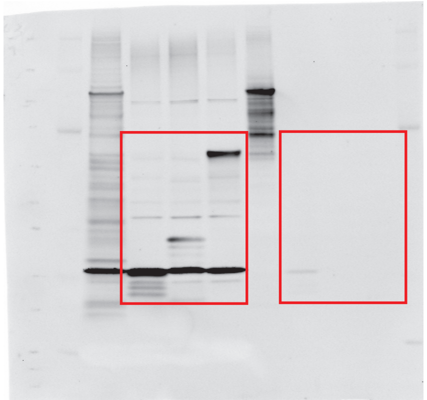

Figure 1F_anti-HA

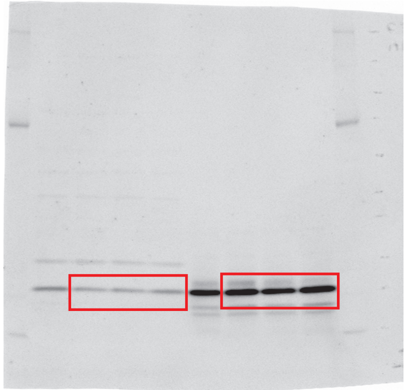

Figure 1-figure supplement1A_CEP89

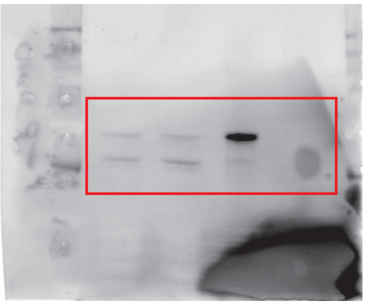

Figure 1-figure supplement1A_NCS1

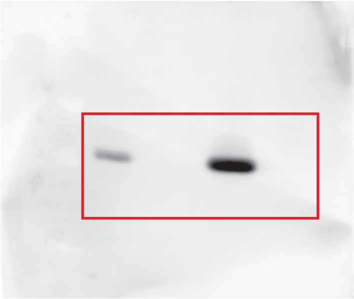

Figure 1-figure supplement 2A_MYC

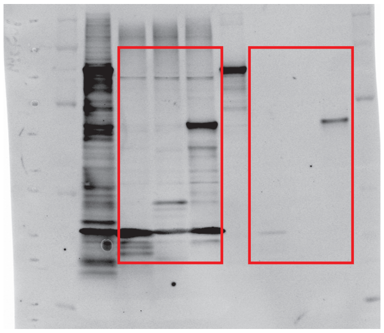

Figure 1-figure supplement 2A_HA

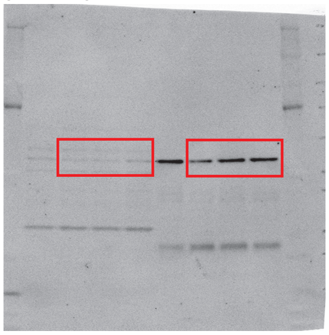

Figure 2-figure supplement 2D_anti-CEP89

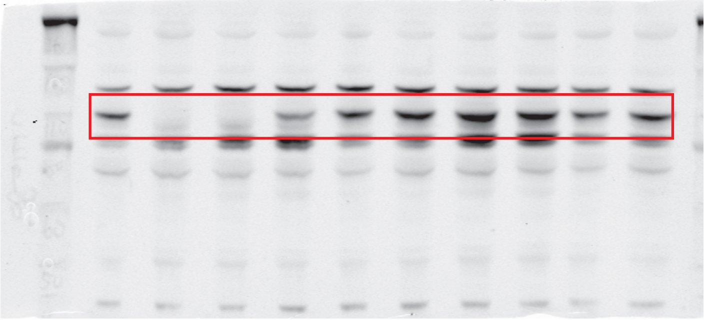

Figure 2-figure supplement 2D_anti-NCS1

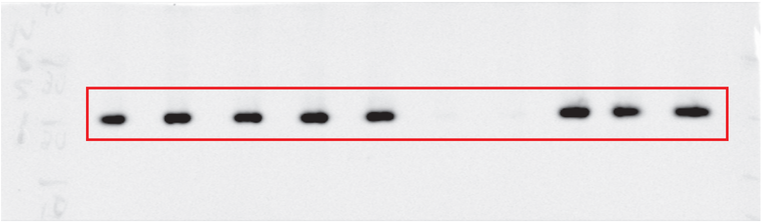

Figure 2-figure supplement 2D_anti-Tubulin

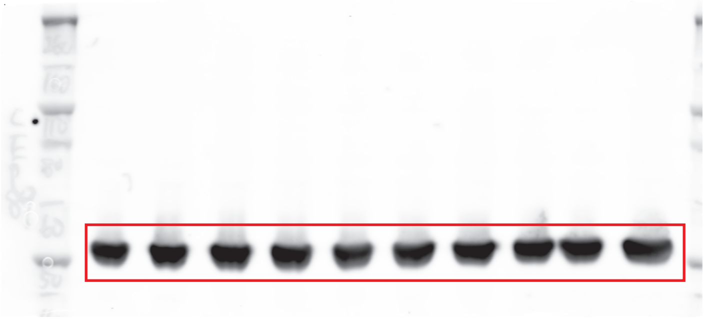

Figure 2-figure supplement 2E_anti-GFP

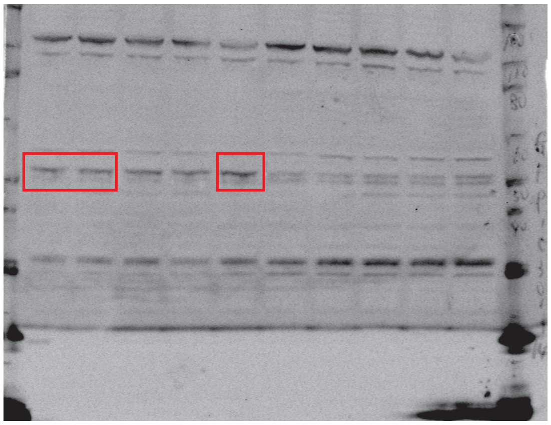

Figure 2-figure supplement 2E_anti-Tubulin

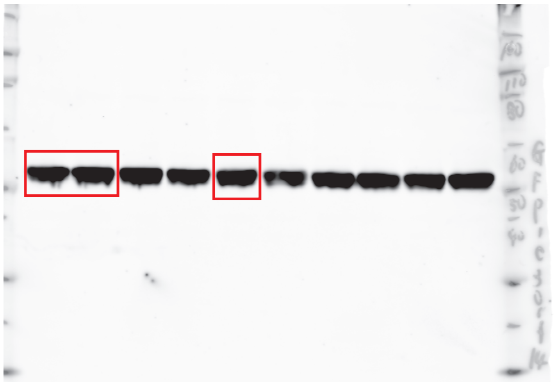

Figure 4A_anti-NCS1

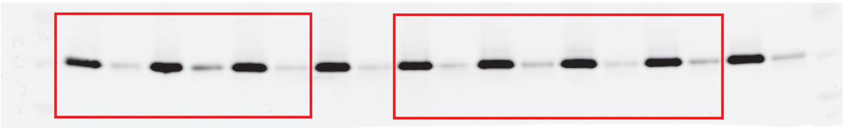

Figure 4A_anti-Tubulin

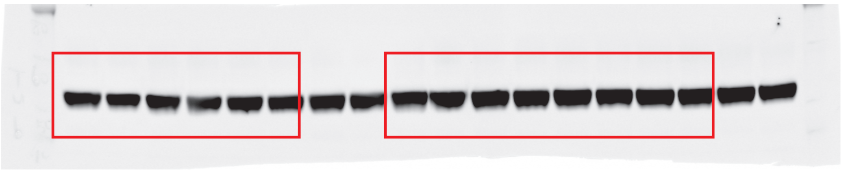

Figure 5A_CEP89

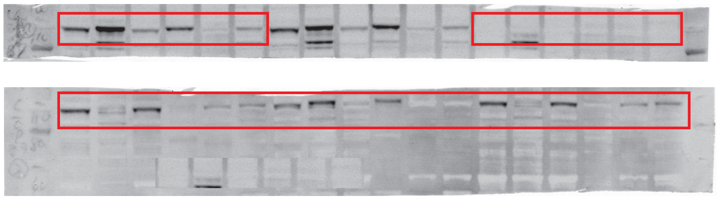

Figure 5A_NCS1

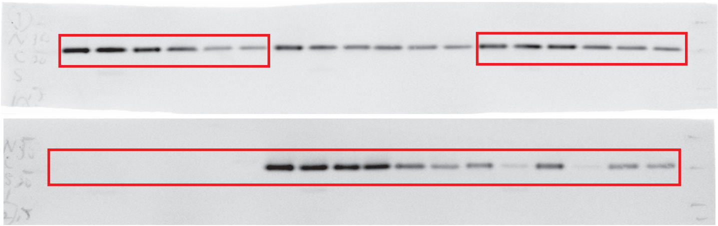

Figure 5A_EGFR

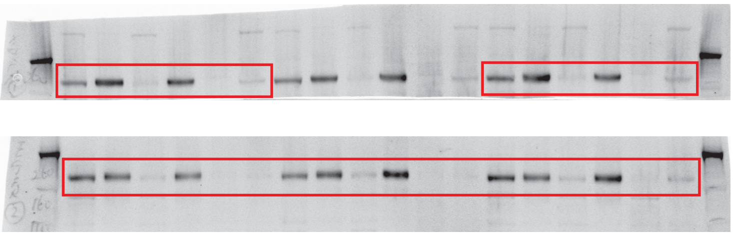

Figure 5A_RabGDI

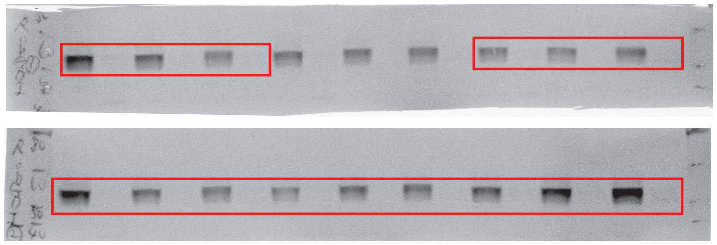

Kanie et al., Resource-full blot No.2

Figure 5B_NCS1

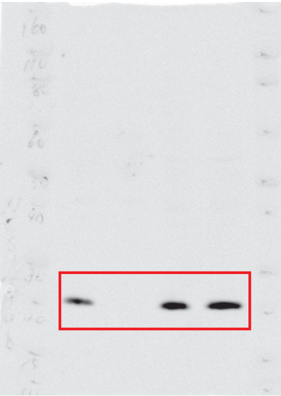

Figure 5B_Tubulin

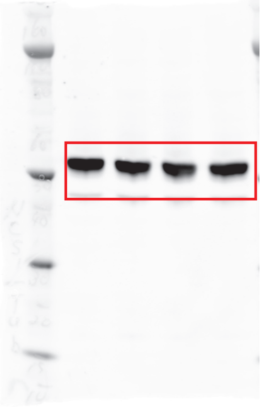

Figure 5-figure supplement 1A_GFP (CEP89)

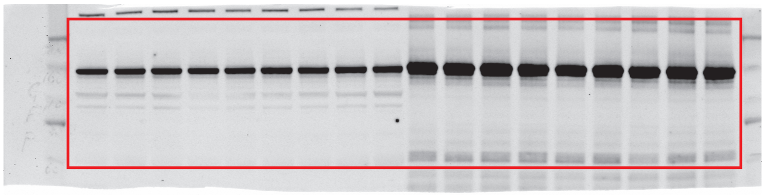

Figure 5-figure supplement 1A_NCS1

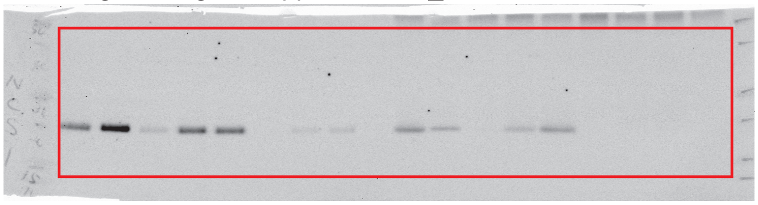

Figure 7-figure supplement 1A_NCS1

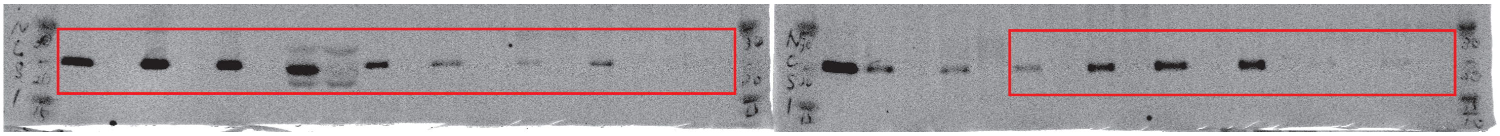

Figure 7-figure supplement 1A_IFT88

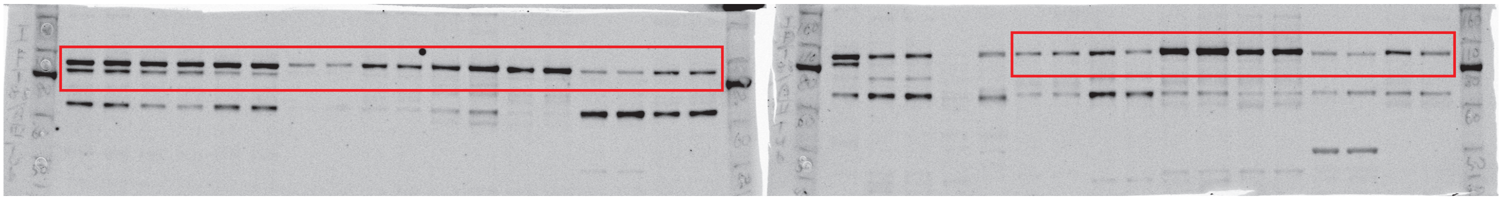

Figure 7-figure supplement 1A_α-Tub

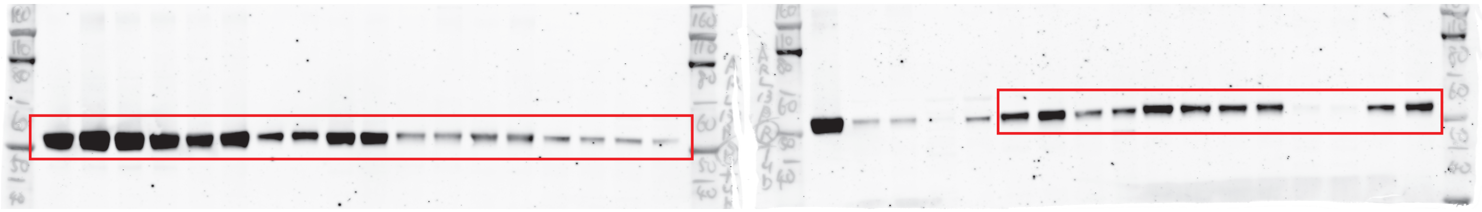

Figure 7-figure supplement 1A_β-TubIII

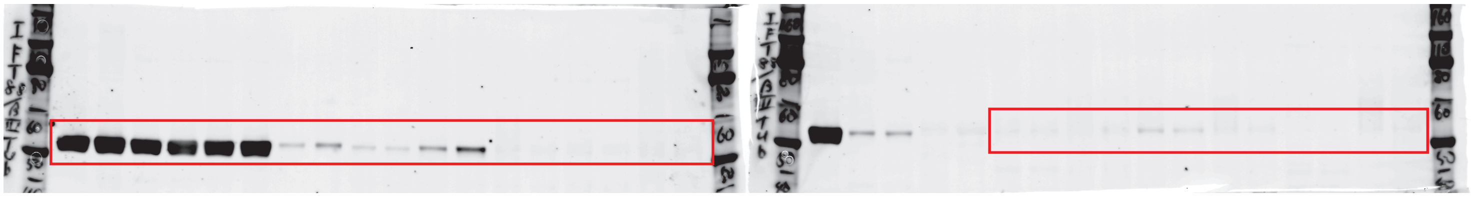
